# TACC3–ch-TOG track the growing tips of microtubules independently of clathrin and Aurora-A phosphorylation

**DOI:** 10.1101/008359

**Authors:** Cristina Gutiérrez-Caballero, Selena G. Burgess, Richard Bayliss, Stephen J. Royle

## Abstract

The interaction between TACC3 (transforming acidic coiled coil protein 3) and the microtubule polymerase ch-TOG (colonic, hepatic tumor overexpressed gene) is evolutionarily conserved. Loading of TACC3–ch-TOG onto mitotic spindle microtubules requires the phosphorylation of TACC3 by Aurora-A kinase and the subsequent interaction of TACC3 with clathrin to form a microtubule-binding surface. Recent work indicates that TACC3 can track the plus-ends of microtubules and modulate microtubule dynamics in non-dividing cells via its interaction with ch-TOG. Whether there is a pool of TACC3–ch-TOG that is independent of clathrin in human cells, and what is the function of this pool, are open questions. Here, we describe the molecular interaction between TACC3 and ch-TOG that permits TACC3 recruitment to the plus-ends of microtubules. This TACC3–ch-TOG pool is independent of EB1, EB3, Aurora-A phosphorylation and binding to clathrin. We also describe the distinct combinatorial subcellular pools of TACC3, ch-TOG and clathrin. TACC3 is often described as a centrosomal protein, but we show that there is no significant population of TACC3 at centrosomes. The delineation of distinct protein pools reveals a simplified view of how these proteins are organized and controlled by post-translational modification.

## Introduction

Microtubules (MTs) are dynamic polymers of α/β-tubulin that are involved in numerous cellular processes, including intracellular transport, chromosome segregation and control of cell shape and migration. Each MT is polarized, having a fast-growing plus-end and a minus-end that grows more slowly. The plus-ends swap between episodes of growth and shrinkage, powered by GTP hydrolysis (Desai and Mitchison, 1997). MT plus-end tracking proteins (+TIPs) bind the plus-ends of MTs, typically during episodes of growth (Akhmanova and Steinmetz, 2010).

Among the best-known +TIPs are the end-binding (EB) proteins (EB1-3) and proteins with TOG or TOG-like domains, e.g. ch-TOG/XMAP215 (Mimori-Kiyosue et al., 2000; van der Vaart et al., 2011). EB proteins can recruit a plethora of proteins, mainly via [ST]X[IL]P motifs, to induce their plus-end tracking behavior (Honnappa et al., 2009). An exception is ch-TOG/XMAP215 which contains no [ST]X[IL]P motifs (Jiang et al., 2012) and tracks MT plus-ends *ahead* of EB proteins (Maurer et al., 2014; Nakamura et al., 2012; Zanic et al., 2013). EB proteins and ch-TOG have different modes of MT binding. EB proteins bind growing ends of MTs, but do not select between plus- and minus-ends, whereas ch-TOG/XMAP215 only binds the plus-ends but does not distinguish between growing and shrinking ends (Maurer et al., 2014; Zanic et al., 2013).

Transforming acidic coiled-coil protein 3 (TACC3) is a cancer-associated protein that binds ch-TOG (Thakur et al., 2013). The interaction is evolutionarily conserved and occurs via a break in the coiled-coil (TACC) domain of TACC3 and a region C-terminal to the TOG6 domain on ch-TOG (Hood et al., 2013; Mortuza et al., 2014; Thakur et al., 2014). The focus on these two proteins has centered on their roles in mitosis. TACC3–ch-TOG complexes were originally proposed to antagonize the function of the MT depolymerase KIF2C/MCAK at spindle poles (Kinoshita et al., 2005) and stimulate MT assembly independently of MCAK (Barr and Gergely, 2008). TACC3 is a substrate of Aurora-A kinase, which phosphorylates TACC3 on serine 558 (Cheeseman et al., 2011; Kinoshita et al., 2005; LeRoy et al., 2007). This phosphorylation permits TACC3 to bind clathrin heavy chain, whereupon the TACC3–clathrin can bind MTs in concert (Booth et al., 2011; Fu et al., 2010; Hood et al., 2013; Hubner et al., 2010; Lin et al., 2010). The ternary complex of TACC3–ch-TOG–clathrin is involved stabilizing kinetochore fibers of the mitotic spindle by inter-MT bridging (Booth et al., 2011). This information, and the observation that removal of clathrin by knocksideways removes all TACC3 from the spindle cast doubt on whether a pool of TACC3–ch-TOG binary complex is present in mitotic cells (Cheeseman et al., 2013).

Recently, Nwagbara et al. described a +TIP function for xTACC3–XMAP215 in non-dividing *Xenopus* cells and reported an effect of xTACC3 on MT dynamics in neuronal growth cones (Nwagbara et al., 2014). This work raised several questions. For example, is TACC3 a +TIP in human cells? Does +TIP activity occur during mitosis and/or interphase? Is it dependent on interaction with clathrin and/or ch-TOG? What is the cell biological function of the TACC3 +TIP activity?

We set out to investigate these issues and define the subcellular locations of each protein, beginning by examining the dynamics of TACC3 on MTs. We discovered that a fraction of TACC3 behaves as a +TIP in human cells, with ch-TOG mediating the association of TACC3 with the MT plus-end. This subcomplex is distinct from the Aurora-A-phosphorylated form of TACC3 that associates with clathrin in the TACC3–ch-TOG–clathrin complex on K-fibers during mitosis. Using this information, we describe the pools of TACC3, ch-TOG and clathrin, alone and in combination at different stages of the cell cycle.

## Materials and Methods

### Cell culture

HeLa cells (HPA/ECACC #93021013) were maintained in Dulbecco’s Modified Eagle’s Medium (DMEM) plus 10% fetal bovine serum (FBS) and 100 U/ml penicillin/streptomycin. HeLa Kyoto cells stably transfected with a BAC to express human TACC3 with a C-terminal GFP tag (Hubner et al., 2010) were a kind gift from Tony Hyman (Dresden). Two lines were used MCP_ky_179 and 190, with identical results and were maintained in the same way as regular HeLa. RPE1 cells stable expressing EB3-tdTomato (Theisen et al., 2012), were a kind gift from Anne Straube (Warwick). RPE1 cells were cultured in DMEM/Nutrient Mixture F-12 Ham (Sigma) supplemented with 10% FBS and 100 U/ml penicillin/streptomycin, 2 mM L-Glutamine and 0.25% Sodium bicarbonate (Sigma). RPE1 stably expressing EB3-tdTomato were maintained in medium containing 500 *µ*g/ml G418. All cells were cultured at 37°C and 5% CO_2_in humidified incubator. HeLa and RPE1 cells were transfected using GeneJuice (Novagen) and FuGENE 6 (Promega) respectively and Lipofectamine2000 (Life Technologies) was used for transfection of siRNA. Transfections were carried out according to manufacturer’s instructions in all cases.

### Molecular Biology

Most plasmids were available from previous work (Booth et al., 2011; Hood et al., 2013; Royle et al., 2005). To deplete endogenous ch-TOG and re-express ch-TOG-GFP, an existing plasmid (pBrain-GFP-ch-TOGKDP-shch-TOG) was modified to move the GFP from the N-terminus of ch-TOG to the C-terminus. This plasmid was subjected to site-directed mutagenesis to introduce LL1939,1942AA mutations. The target sequence for TACC3 siRNA was cagtttggaacttcctcgt (SASI_Hs01_00156991 MISSION pre-designed siRNA). Constructs for recombinant expression of TACC3(629-838) and ch-TOG truncates were produced as previously described (Hood et al., 2013). Point mutations were introduced in pETM6T1 ch-TOG(1517-1957) by the Quikchange procedure (Stratagene). For depletion of EB1 and EB3, two siRNAs were used that targeted TGCTCCAGCTCTGAATAAA and ACTATGATGGAAAGGATTAC, respectively (Komarova et al., 2005).

### Microscopy and analysis

For live cell imaging, cells were plated in 35 mm glass bottom fluorodishes. After 48 h after transfection cells were placed in a 37°C chamber on the microscope stage of a spinning disc confocal system (Ultraview Vox, Perkin Elmer) cultured in the CO2-independent media Leibovitz’s L-15 Medium (Sigma) supplemented as for growth conditions. Imaging was performed using a 100X ~1.4 NA oil immersion objective lens. Cells were typically imaged every second for 1 minute. Cells were excited at 488 nm and 561 nm and images captured simultaneously with two cameras (Hamamatsu C10600-10B ORCA-R2). On each day of imaging, camera alignment was performed using 0.5 *µ*m diameter fluorescent beads.

To analyse the intensity of EB3-tdTomato and GFP-TACC3 a 3 pixel-wide line was drawn over the MT end and the intensities for both channels exported. The intensity of each profile was normalised so that the minimum value was zero and the maximum value was 1 (∆F/Fmax). The position of maximum intensity in the EB3 channel was used to offset the EB3 and TACC3 traces, such that this position was 0.

For knocksideways experiments, HeLa cells were transfected with pBrain-GFP-FKBP-TACC3KDP-shTACC3 (depletes endogenous TACC3 and re-expresses tagged TACC3) and pMito-PAGFP-FRB (‘invisible’ MitoTrap). Cells were treated with rapamycin (R8781, Sigma Aldrich) at 200 nM or DMSO 0.1% (control) for 20 min at 37°C. Cells were fixed in methanol at -20°C for 5 min, before immunostaining with mouse anti-EB1 (1:500, BD Transduction Laboratories, 610534) or rabbit anti-ch-TOG (1:5000, Autogen Bioclear, 34032) and Alexa568 conjugated secondary antibodies. Images were taken on a Nikon Ti epifluorescence microscope with 60X oil immersion objective (1.4 NA) and a Hamamatsu Orca-ER camera.

For immunofluorescence, cells were fixed with PTEMF (50 mM Pipes, pH 7.2, 10 mM EGTA, 1mM MgCl_2_, 0.2% Triton X-100, and 4% paraformaldehyde) for 15 min at RT, and then permeabilized (PBS/0.1% Triton X-100) for 10 min. Cells were blocked (PBS, 3% BSA, and 5% goat serum) for 1 h, and then incubated for 1 h with the specified primary antibodies: mouse anti-CHC (X22, 1:1000), mouse anti-TACC3 (AbCam ab56595, 1:1000), rabbit anti-Pericentrin (Abcam ab4448, 1:5000), mouse anti-CENP-A (Abcam, ab13939, 1:500). Secondary antibodies, Alexa Fluor 488 and 568 (Life Technologies, 1:500). Cells were rinsed with PBS and mounted with mowiol containing DAPI. Images were taken using a spinning disk confocal with a 60X objective and a z-step of 0.5 *µ*m.

For migration analysis, RPE1 cells were seeded into Lab-Tek dishes (Nunc) coated with 10 μg/ml fibronectin (Sigma). On the day of imaging, nuclei were stained with NucRed^™^ Live 647 (R37106, Life technologies) and then imaged for 6 h at a rate of one image every 6 min or 20 min. Imaging was done using a Nikon Ti microscope and Hamamatsu Orca-ER camera, using a 20X air objective. Cell movements were tracked using the Manual Tracking plug-in in Fiji/ImageJ to monitor the xy position of the centre of the nucleus. 2D Coordinates were fed to IgorPro for further analysis using custom-written procedures. A code snippet for rotation of a 2D set of co-ordinates about the origin is deposited at http://www.igorexchange.com/node/5895

MT dynamics were analyzed from movies (1 Hz) of EB3-tdTomato RPE1 cells using uTrack 2.1.0 in MATLAB R2013b (Applegate et al., 2011). The following parameters were used for all movies. Anisotropic Gaussian Detection method; Maximum Gap length 4 frames; Minimum length of Track Segments 3 frames; Brownian Search Radius 2-10 px (Multiplication Factor 3); Number of Frames for Nearest Neighbor Distance Calculation 5; Maximum forward angle 30°; Maximum backward angle 10°; Maximum shrinkage factor relative to growth speed 1.5; Fluctuation radius 1 px.

Kymographs and temporal color-coded images of movies were generated in ImageJ. Kymographs were contrast adjusted for Images were cropped in ImageJ or Photoshop and figures were assembled in Illustrator CS5.1. Box plots show the median, 75^th^ and 25^th^ percentile and whiskers show 90^th^ and 10^th^ percentile.

### Biochemistry

Western blotting was as described previously (Kaur et al., 2014). Using the following primary antibodies: mouse anti-EB1 (BD Transduction labs, 610534, 1:2000), mouse anti-EB3 (BD Transduction labs, 612156, 1:1000), mouse anti-alpha tubulin (DM1A Abcam ab7291, 1:10000), mouse anti-TACC3 (Abcam ab56595, 1:500). A rabbit polyclonal antibody was raised against peptides, EVIEGYRKNEESLKKC and TVEQKTKENEELTRIC to recognize the TACC domain of TACC3 (Eurogentec). Expression and purification of recombinant TACC3(629-838) and ch-TOG truncates was carried out as stated in earlier work (Hood et al., 2013). *In vitro* binding assays between TACC3(629-838) and ch-TOG truncates were performed as previously stated (Hood et al., 2013). For CD spectroscopy, wild type and point mutants of ch-TOG(1517-1957) were dialyzed into 20 mM sodium phosphate pH 7.0, 50 mM NaCl and diluted to 0.1 mg/ml. Spectra were collected on a Chirascan+ instrument (Applied Photophysics) using a 0.01 cm pathlength quartz cell at 20°C and are shown as the average of three replicates after baseline subtraction and smoothing. ClustalW2 was used to align ch-TOG sequences (Larkin et al., 2007).

## Results

### TACC3 is a microtubule plus-end tracking protein in human cells

To investigate the potential +TIP activity of TACC3, we used live-cell spinning disk confocal microscopy of human cell lines expressing GFP-tagged TACC3. In interphase and mitotic cells, GFP-TACC3 formed clear punctate structures that moved in a directed manner, suggesting that TACC3 could behave as a MT plus-end tracking protein (+TIP). Figure 1 shows examples of TACC3 +TIP behavior (see also Movies 1-3). GFP-TACC3 was transiently expressed in retinal pigment epithelial cells (RPE1) stably expressing the end-binding protein EB3 tagged with tdTomato (Fig 1A). In this cell line, EB3-tdTomato tracked the growth of MT plus-ends in a comet-like manner as previously described (Akhmanova and Steinmetz, 2010; Nakagawa et al., 2000). Plus-end tracking of GFP-TACC3 was clearest in interphase where long periods of MT growth were marked by a small punctum of GFP-TACC3 fluorescence. However, in contrast to EB3-tdTomato, GFP-TACC3 stayed at the tip of MTs as they underwent shrinkage (Fig 1A, arrow). In mitotic cells, the +TIP behavior of TACC3 was most clear on the astral MTs (Fig 1A). At metaphase, these signals were often difficult to discern against the fluorescence on the K-fibers of the spindle. Plus-end tracking of TACC3 was much clearer at anaphase on astral and interpolar MTs in either HeLa or RPE1 cells (see below).

**Figure 1.**
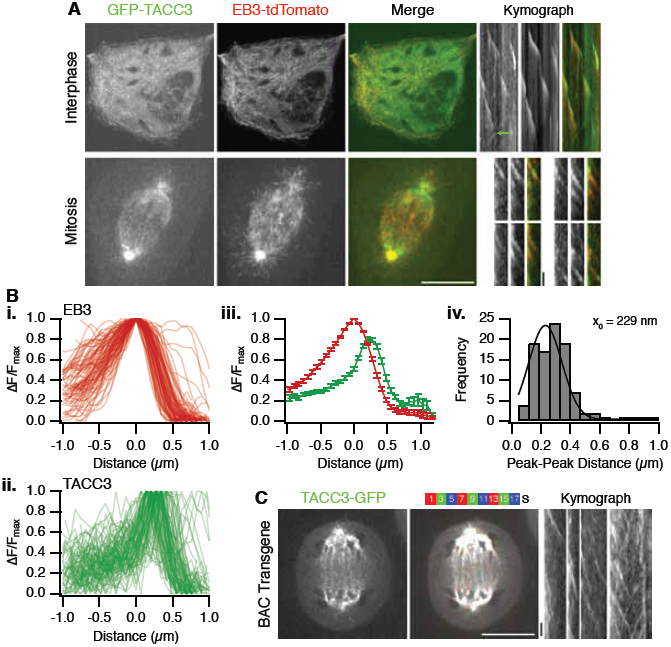
TACC3 is a +TIP that tracks plus-ends of microtubules ahead of EB3. **(A)** Single frames from live-cell imaging experiments of RPE1 cells expressing GFP-TACC3 (green) in interphase and mitosis. RPE1 cells were stably expressing EB3-tdTomato (red in merge). Similar results were seen in the parental RPE1 cell line. For each condition, a single frame from the movie (1 Hz) is shown together with contast-enhanced kymographs to illustrate the movement of TACC3 puncta over time. A representative MT shrinkage event is marked by a green arrow. **(B)** Plot of fluorescence intensity as a function of distance at growing MT ends for (i) EB3-tdTomato, ii) GFP-TACC3 in interphase RPE1 cells. The change in fluorescence intensity was normalized to the brightest point in the profile and the distance for both channels set to 0 at the position of brightest intensity of EB3. Mean ± s.e.m. of all profiles (iii). Histogram of the distance from EB3 peak to TACC3 peak (iv). A single Gaussian fit is shown with a peak at 229 nm. Mean peak-peak distance was 284 nm. N=100. **(C)** Single frame (left) and color projection of live cell images of HeLa Kyoto cells with stably integrated BAC transgene to express TACC3-GFP under its native promoter. Late anaphase cells were imaged at 0.5 Hz and nine consecutive frames were projected into one image using different colors as indicated. Numbers indicate time (s). Scale bar, 10 μm (20 μm for interphase) and 10 s.

In interphase or mitotic cells, GFP-TACC3 was present at the very distal tip of the growing MT, apparently travelling ahead of the EB3 signal (Fig 1A, kymographs). To analyze this more rigorously, fluorescence intensities of EB3-tdTomato and GFP-TACC3 were extracted as a function of distance along a growing MT track (see Methods). On average, the peak GFP-TACC3 intensity was 229 nm ahead of the peak EB3-tdTomato signal (Fig 1B). This finding agrees with work on xTACC3 and also its binding partner XMAP215/ch-TOG (Maurer et al., 2014; Nakamura et al., 2012; Nwagbara et al., 2014).

Was the +TIP activity of TACC3 an artifact of transient expression of GFP-TACC3? Although we only imaged cells expressing very low levels of GFP-TACC3 – because overexpression resulted in aggregation of TACC3 (Gergely et al., 2000a) and no discernable +TIP activity – we wanted to rule out the possibility that +TIP behavior was an artifact of over-expression. As an alternative, we imaged live HeLa Kyoto cells expressing TACC3-GFP from a BAC transgene (Hubner et al., 2010). This protein is expressed at close-to-endogenous levels by virtue of the endogenous promoter. Again, TACC3 tracking the growth of MT plus-ends could clearly be seen (Fig 1C). Together these experiments confirm that TACC3 is a +TIP in human cells in interphase or mitosis and that it tracks the very distal tip of growing MTs.

### TACC3 +TIP behavior is independent of EB1 and EB3

End-binding proteins, such as EB1 and EB3, recruit proteins to the growing tips of MTs by virtue of [ST]X[IL]P motifs. Although TACC3 has no [ST]X[IL]P motif, it was detected in a pull-down for GST-EB1 binding proteins, albeit with low confidence (Jiang et al., 2012). To test whether or not EB proteins play a role in TACC3 +TIP behavior, we visualized GFP-TACC3 in cells depleted of EB1 and EB3. We targeted these two end-binding proteins because EB1 and EB3, but not EB2 promote persistent microtubule growth, and also knockdown of EB1-3 is not possible (Komarova et al., 2009). GFP-TACC3 +TIP tracking was indistinguishable from cells transfected with control siRNA (Fig 2A). We confirmed that good depletion of both EB1 and EB3 was achieved in these experiments (Fig 2B). These experiments indicated that EB1 and EB3 are not required for TACC3 to bind to MT plus-ends.

**Figure 2.**
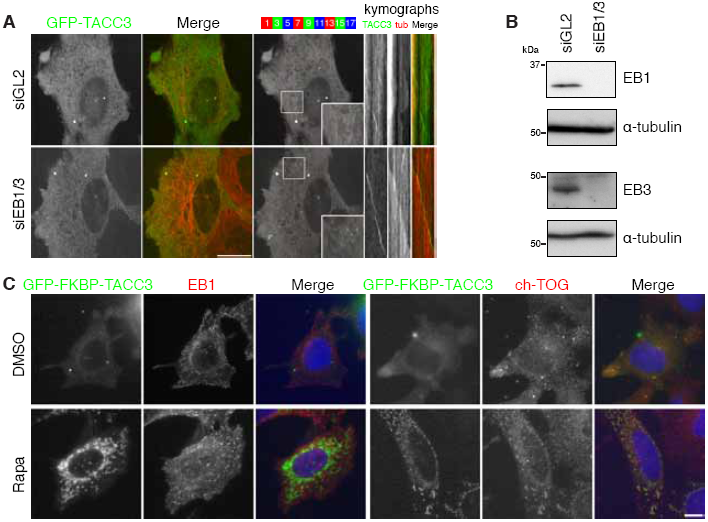
TACC3 +TIP activity is independent of EB1 and EB3. **(A)** Single frame, merge and color projection of live-cell imaging of HeLa cells expressing mCherry-tubulin (red) and GFP-TACC3 (green) that were transfected with either control siRNA (siGL2) or two siRNAs targeting EB1 and EB3 (siEB1/3). Cells in interphase together with typical kymographs are shown (right). Scale bar, 20 *µ*m and 10 s. **(B)** Western blots to show the amount of EB1 or EB3 remaining following double depletion of EB1 and EB3 from cells used for imaging in B. Blotting for alpha-tubulin was used as a loading control. **(C)** Representative micrographs of a knocksideways experiment to test for co-rerouting of EB1 or ch-TOG with TACC3. HeLa cells expressing GFP-FKBP-TACC3 and PAGFP-MitoTrap (not shown) were treated as indicated and fixed with methanol before staining with anti-EB1 or anti-ch-TOG. Scale bar, 10 *µ*m.

Next, we wanted to use a similar approach to test the involvement of ch-TOG in TACC3 localization at MT plus-ends. However, in cells transfected with siRNA (or shRNA) targeting ch-TOG, GFP-TACC3 still acted as a +TIP. This turned out to be due to inefficient depletion of ch-TOG in cells expressing GFP-TACC3 (data not shown), as reported elsewhere (Nwagbara et al., 2014).

As an alternative, we used the knocksideways system (Cheeseman et al., 2013) to test for an interaction *in situ* between TACC3 and endogenous ch-TOG in interphase cells. TACC3-depleted HeLa cells expressing GFP-FKBP-TACC3 and PAGFP-MitoTrap were treated with rapamycin (200 nM) or DMSO (0.1%) as a control. Fig 2C shows that GFP-FKBP-TACC3 was rerouted to mitochondria upon addition of rapamycin. Endogenous EB1, detected by immunofluorescence remained in comets despite the rerouting of TACC3 to mitochondria, consistent with a lack of interaction between TACC3 and EB1. By contrast, endogenous ch-TOG was co-rerouted with GFP-FKBP-TACC3 to mitochondria (Fig 2C), verifying that the interaction between TACC3 and ch-TOG occurs during interphase. These observations are consistent with a model where TACC3–ch-TOG binds the plus-ends of MTs independently of EB1.

### TACC3 +TIP behavior depends on its interaction with ch-TOG and is independent of Aurora-A phosphorylation and subsequent interaction with clathrin

To determine if TACC3 +TIP behavior was indeed dependent on ch-TOG, we expressed GFP-tagged TACC3 deletion mutants that had previously been shown to be unable to bind ch-TOG (Hood et al., 2013). No +TIP activity was seen at interphase or anaphase for two different mutants, ∆678-681 or ∆682-688. The lack of plus-end tracking is not due to protein misfolding because, we have previously shown that either deletion does not interfere with protein structure, and both mutants were still able to localize to the mitotic spindle (Fig 3) (Hood et al., 2013). These results indicate that TACC3–ch-TOG complexes can track the plus-ends of MTs and that plus-end recognition is by ch-TOG and not by TACC3.

**Figure 3.**
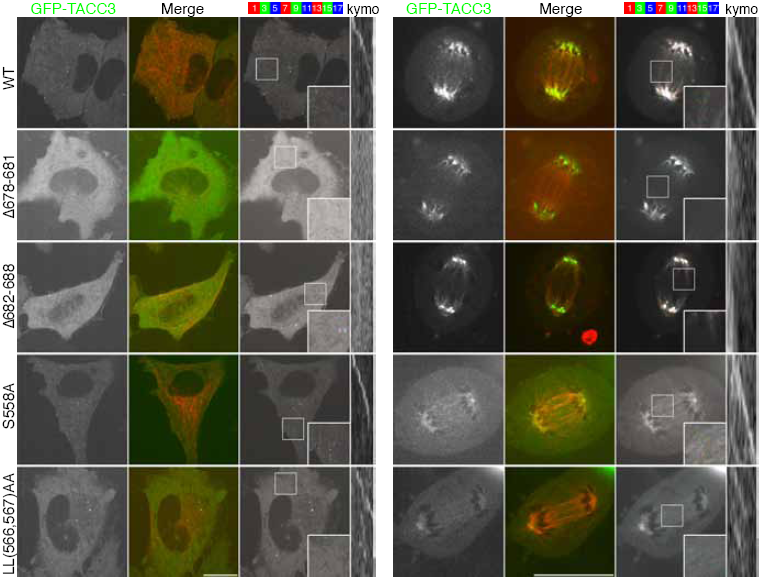
TACC3 +TIP activity is via its interaction with ch-TOG and is separate from the TACC3–ch-TOG–clathrin inter-MT bridge complex. Single frame, merge and color projection of live-cell imaging of TACC3-depleted HeLa cells expressing mCherry-tubulin (red) and RNAi-resistant GFP-TACC3 (green), mutants that do not interact with ch-TOG (∆678-681, ∆682-688), or two mutants that cannot bind clathrin: non-phosphorylatable mutant (S558A) or dileucine mutant (LL566,567AA). Cells in interphase or anaphase together with typical kymographs are shown (right). Similar results were achieved in RPE1 cells. Scale bar, 20 *µ*m and 10 s.

We next tested the hypothesis that TACC3–ch-TOG +TIP activity was separate from the TACC3–ch-TOG–clathrin inter-MT bridge complex. Formation of the TACC3–ch-TOG–clathrin complex depends on Aurora-A phosphorylation of S558 on TACC3, allowing a dileucine motif (566,567) to bind to the ankle of clathrin heavy chain. We previously showed that non-phosphorylatable TACC3(S558A) and a TACC3 mutant in which the dileucine motif had been mutated (LL566,567AA) were unable to bind clathrin and could not localize to the mitotic spindle (Hood et al., 2013). Live-cell imaging of either of these mutants in HeLa cells depleted of endogenous TACC3 showed +TIP tracking comparable to wild type GFP-TACC3 in interphase and anaphase (Fig 3). Moreover, since +TIP activity for TACC3 was observed in interphase, a time when 1) Aurora-A activity is low and 2) clathrin is not known to interact with TACC3, the +TIP activity of TACC3 is independent of clathrin-binding and phosphorylation by Aurora-A.

Can ch-TOG track MT plus-ends independently of TACC3? To address this question, we sought a ch-TOG mutant that cannot bind to the TACC domain of TACC3 and tested whether this mutant can track MT plus-ends. To do this, we first narrowed down the TACC3-binding region within ch-TOG by testing for co-precipitation of recombinant ch-TOG fragments with a His-NusA tagged TACC3 fragment (629-838) using His-NusA as a control. Binding of ch-TOG(1517-1957) but not of ch-TOG(1517-1932) was observed, suggesting that the region of ch-TOG that mediates binding is residues 1932-1957 (Fig 4A). Alignment of these regions from ch-TOG orthologs highlighted a conserved patch including a pair of leucines which we targeted for mutation (Fig 4B). Mutation of either L1939 or L1942 to either alanine or arginine blocked the ability of ch-TOG(1517-1957) to co-precipitate TACC3(629-838) (Fig 4C). Circular dichroism spectra of the L1939A and L1942A mutant proteins were similar to that of wild type ch-TOG(1517-1957) suggesting that these mutations had no significant effect on folding (Fig 4D).

**Figure 4.**
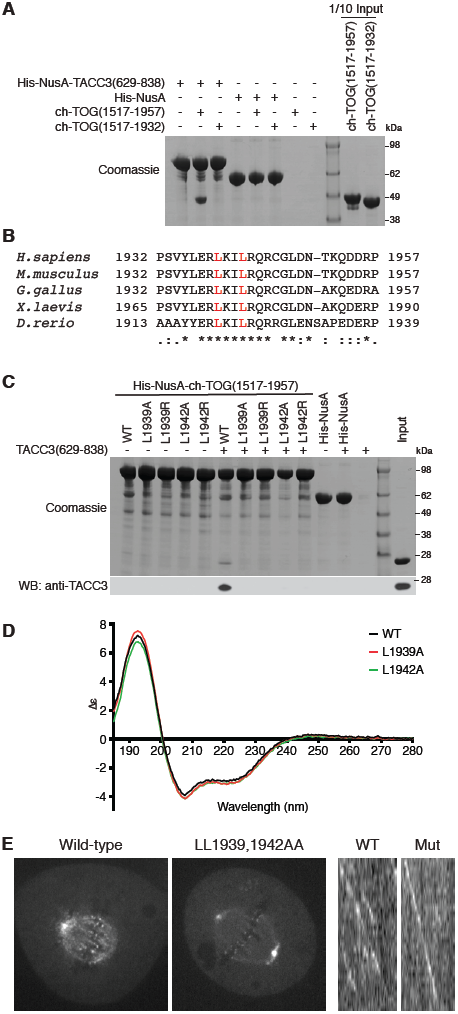
A ch-TOG mutant deficient in TACC binding localizes to centrosomes and kinetochores and tracks the plus-ends of MTs. **(A)** Co-precipitation assay between His-NusA-TACC3(629-838) and fragments of the C-terminal region of ch-TOG. His-NusA-TACC3(629-838) was bound to Ni Sepharose beads and incubated with ch-TOG proteins. His-NusA was used as a tag binding control. **(B)** Sequence alignment of the TACC3-binding region of ch-TOG orthologs. Identical residues are represented with ‘*’, conserved amino acids with ‘:’ and semi-conserved residues with ‘.’. Leucine residues marked in red were targeted for mutation. **(C)** Co-precipitation assay between wild type and point mutants of His-NusA-ch-TOG(1517-1957) and TACC3(629-838). His-NusA-ch-TOG(1517-1957) wild type and mutant proteins were bound to Ni Sepharose beads and then incubated with TACC3(629-838). His-NusA was used as a tag binding control. The reactions were separated by SDS-PAGE (above) and subject to Western blot using an anti-TACC3 TACC domain antibody (below). Input was 1/10 of the binding reaction for Coomassie and 1/50 for Western blot. **(D)** Circular dichroism spectroscopy of wild type ch-TOG(1517-1957) and L1939A and L1942A mutants. **(E)** Confocal micrographs of ch-TOG-depleted HeLa cells expressing ch-TOG-GFP wild type (left) or LL1939,1942AA double mutant (centre). Kymographs showing similar interphase plus-end tracking for wild type (WT) and LL1939,1942AA double mutant (Mut) ch-TOG-GFP (right).

Having identified mutations that knock-out the interaction with TACC3, we next tested the effect of these mutations on ch-TOG-GFP +TIP behavior. A ch-TOG-GFP(LL1939,1942AA) construct was expressed in ch-TOG-depleted HeLa cells and its subcellular distribution studied by confocal microscopy. Wild type ch-TOG-GFP was localized to kinetochores, spindle MTs and centrosomes (Fig 4E), whereas the mutant was not associated with spindle MTs as expected, but still localized to the centrosomes and kinetochores. We return to these observations later. Live cell imaging showed that the mutant tracked the plus-ends of MTs in interphase similarly to the wild-type (Fig 4F). All cells imaged showed clear MT plus-end tracking (n>20), confirming that +TIP tracking of ch-TOG does not require TACC3.

### A potential role for +TIP activity of TACC3 in interphase cell migration?

What is the cellular function of TACC3 binding to ch-TOG at the plus-ends of MTs? Analysis of this novel population of TACC3 was not possible during mitosis, because depleting or mutating TACC3 interferes with its role in stabilizing K-fibers as part of the TACC3–ch-TOG–clathrin complex. During interphase however, this complex is not formed and no mitotic spindle is present; we therefore reasoned that this stage of the cell cycle was the best time to investigate the cellular function of TACC3 +TIP activity. Previous work suggested that TACC3 may influence cell migration (Ha et al., 2013). If this is correct, then any changes in cell migration might be due to altered MT dynamics. We began by testing a role for TACC3 in interphase cell migration.

RPE1 cells transfected with siRNAs targeting GL2 (control) or TACC3 were plated on fibronectin-coated dishes and imaged over 6 h and their 2D migration observed (see Methods). TACC3-depleted cells moved more slowly than control cells and the cumulative distance that they migrated was on average less than the control population (Fig 5A, 5B). Aligning the individual migration tracks so that their end position was along the same axis, gave the impression that growth was more directed after TACC3 depletion, and that the cells made fewer turns (Fig 5C). However, this apparent effect could be explained by the lower migration speed in TACC3-depleted cells because the directionality ratio between distance migrated and displacement (d/D) was not significantly different from the control group (Fig 5D). Mean squared displacement (MSD) analysis also showed that the directional persistence was similar between TACC3-depleted cells and control cells and that differences in migration were due to a lower migration speed (Fig 5E). Over three experiments, the migration speed for TACC3-depleted cells was ~30% lower than control cells (Fig 5F). These results indicate that TACC3 is involved in cell migration and they open the possibility the +TIP behavior of TACC3 may underlie this phenotype.

**Figure 5.**
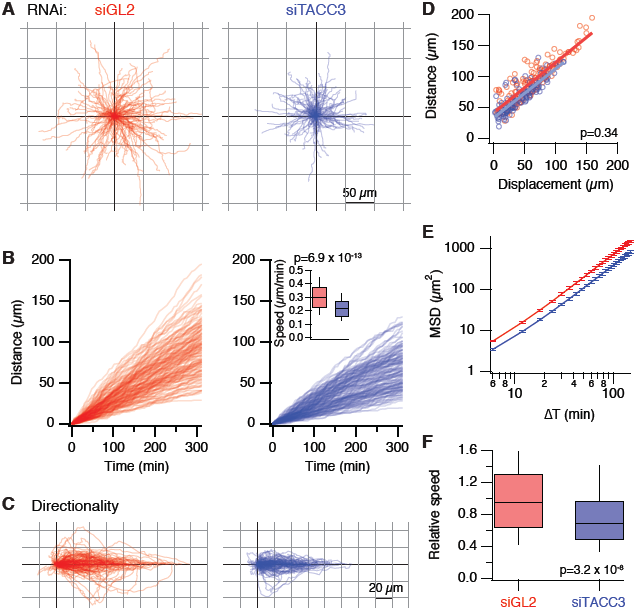
Role for TACC3 in interphase cell migration. **(A)** Overlay of individual tracks (nuclear position over time) of RPE1 cells transfected with siRNA targeting GL2 (siGL2, red) or TACC3 (siTACC3, blue). Tracks were aligned to the origin. **(B)** Plots to show cumulative distance as a function of time for the tracks shown in A. N_cell_= 147 (GL2), 154 (TACC3), from a single experiment. Inset: Box plot to show the speed of migration for the tracks shown. **(C)** Overlay of individual tracks to assess directionality. Tracks from A were rotated so that the end point of each track aligned with the origin along the x-axis. **(D)** Plot of migration distance as a function of displacement. Displacement is the Euclidian distance from the origin to the endpoint. Gradients of the lines of best fit (commonly referred to as directionality ratio d/D) were similar, 0.81 (siGL2) and 0.79 (siTACC3). **(E)** Plot of mean squared displacement (MSD) analysis of the tracks for lag times ranging from 6 s to 150 s. Errors represent s.e.m. **(F)** Box plot to show the relative speed of TACC3-depleted RPE1 cells versus control cells over three independent experiments. P values derived from Student’s t-test are shown in B, D and F.

### Altered TACC3 expression has no detectable effect on MT dynamics

To test for changes in MT dynamics we used live cell imaging of RPE1 cells stably expressing EB3-tdTomato. Cells were transfected with control siRNA (siGL2) or siRNA targeting TACC3, or plasmids to express GFP or GFP-TACC3 (TACC3 overexpression). MT dynamics were measured using automated particle tracking analysis (see Methods). We found that neither depletion nor overexpression of TACC3 altered MT dynamics relative to their respective control (Fig 6A). The length of EB3 tracks measured, their lifetimes and their resultant speed was unaltered relative to the control (Fig 6A). Analysis of TACC3 depletion in cells used for imaging experiments showed good depletion (Fig 6B). We conclude that the +TIP behavior of TACC3 does not detectably influence MT dynamics in interphase RPE1 cells. Therefore, the changes in cell migration are unlikely to be due to altered MT dynamics.

**Figure 6.**
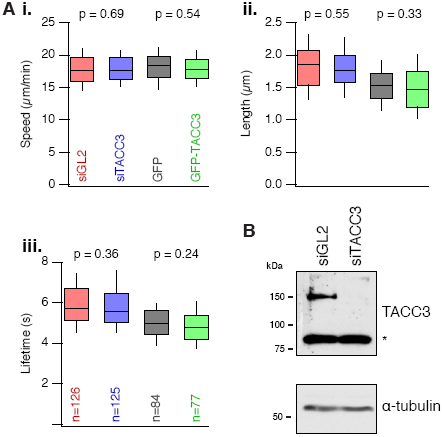
TACC3 does not detectably influence MT dynamics. **(A)** Box plots to show the speed (i), length (ii) and lifetime (iii) of growing MT tracks in interphase RPE1 cells stably expressing EB3. Cells were transfected with either control siRNA (siGL2), siRNA targeting TACC3 (siTACC3), or plasmids to express GFP or GFP-TACC3. MT dynamics were measured from 77-126 cells from three separate experiments. **(B)** Western blots to show the amount of TACC3 remaining following RNAi of RPE1 cells used for imaging. TACC3 runs at ~150 kDa, asterisk indicates a non-specific band. Blotting for alpha-tubulin was used as a loading control.

### Distinct pools of TACC3, ch-TOG, clathrin

The +TIP pool of TACC3–ch-TOG and the observation of ch-TOG targeting to centrosomes and kinetochores independently of TACC3, prompted us to re-examine the subcellular distributions of TACC3, ch-TOG and clathrin. The aim was to document the possible combinatorial pools of TACC3, ch-TOG and clathrin in cells in mitosis or interphase. In mitotic cells, besides the +TIP pool of TACC3–ch-TOG described above, endogenous TACC3 detected by immunofluorescence is present on the K-fibers together with clathrin and ch-TOG, but is largely absent from centrosomes. In addition to localizing to K-fibers, clathrin is found in coated pits and vesicles, while ch-TOG-GFP is also located at the centrosomes and kinetochores (Fig 7A). In interphase cells, ch-TOG is again found at centrosomes while TACC3 is absent from the centrosome in the majority of cells (Fig 7A, 7B). In cells where TACC3 was in the vicinity of pericentrin staining, the two signals did not overlap and TACC3 might be present on MTs here rather than the centrosomes themselves (Fig 7C).

**Figure 7.**
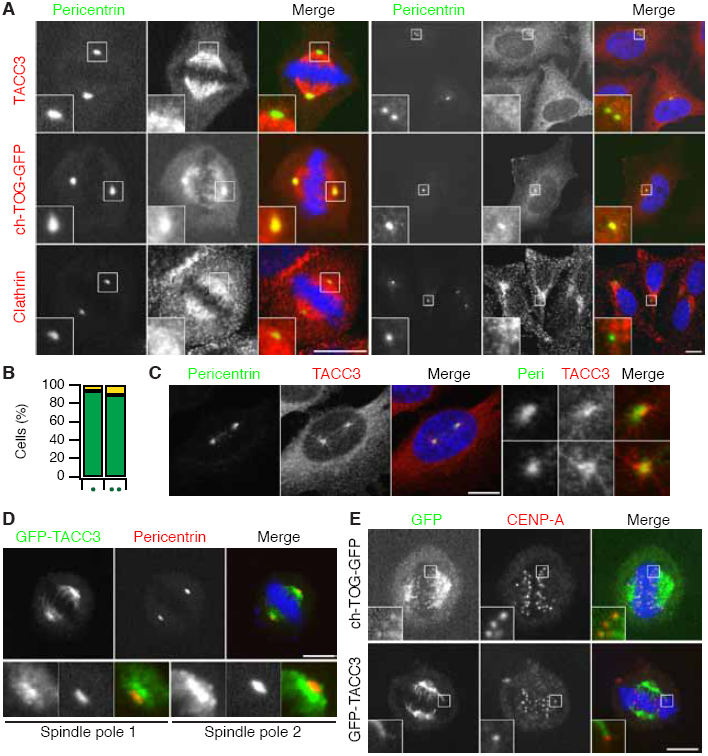
Distinct combinatorial pools of TACC3, ch-TOG and clathrin. **(A)** Typical confocal micrographs to show the subcellular distributions of TACC3, ch-TOG and clathrin in mitotic (left) and interphase (right) HeLa cells. Untransfected cells were fixed and stained for pericentrin and either TACC3 or clathrin heavy chain, or in the case of ch-TOG, were transfected to express ch-TOG-GFP on a background of ch-TOG depletion and then fixed and stained for pericentrin (green in merge). Note the lack of colocalization of TACC3 and pericentrin in surrounding cells. For mitotic cells a single plane of a z-stack is shown. For interphase cells, a maximum intensity z-projection is shown. Zoomed regions show comparable magnification (2X and 4X for mitotic and interphase respectively). Scale bar, 10 *µ*m. **(B)** Bar chart to show the percentage of interphase cells with TACC3 in the vicinity of pericentrin staining (yellow) or no detectable enrichment (green). Results are shown for cells with one or two pericentrin puncta as indicated. **(C)** Single confocal image to show an interphase cell that had TACC3 in the vicinity of centrosomes. Right, 3X enlargements of the centrosomal regions. **(D)** Single confocal image to show a mitotic HeLa cell expressing GFP-TACC3 (green) co-stained for pericentrin (red). Below, 3X enlargements of the centrosomal regions. **(E)** Single confocal image of mitotic HeLa cells expressing ch-TOG-GFP or GFP-TACC3 (green) co-stained for CENP-A. Insets, 3X enlargements of the indicated regions. Scale bars, 10 *µ*m.

The centrosome and kinetochore localization of ch-TOG-GFP was intriguing: why doesn’t its binding partner, TACC3 also localize there? We wondered whether it was an antibody accessibility problem that prevented us from detecting TACC3 at either location. Using GFP-TACC3 we again found no co-localization with pericentrin in mitotic cells (Fig 7D). Moreover, GFP-TACC3 was not found at kinetochores, unlike ch-TOG-GFP (Fig 7E). Finally, recall that ch-TOG-GFP(LL1939,1942AA), a mutant that cannot bind TACC3, was localized at centrosomes and kinetochores but not at the K-fibers (Fig 4E), suggesting that this pool of ch-TOG is not associated with TACC3. These localization data suggest that there are four distinct pools of TACC3, ch-TOG and clathrin in mitotic cells and three pools in interphase (Fig 8).

**Figure 8.**
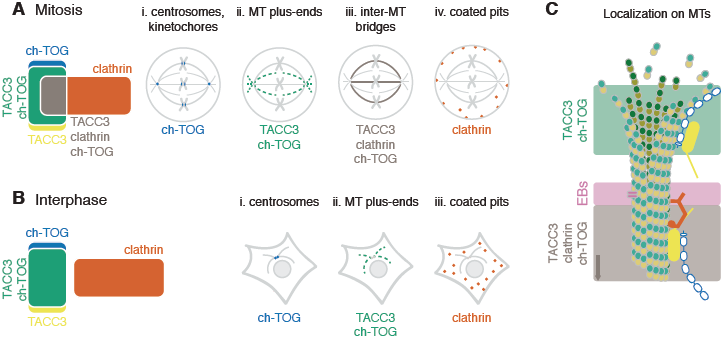
Summary schematic view of distinct combinatorial pools of TACC3, ch-TOG and clathrin. **(A, B)** The cellular pools of TACC3, ch-TOG and clathrin alone or in their possible combinations are shown for a model cell in mitosis (A) or interphase (B). Cytoplasmic populations of each protein can exchange with the corresponding pools and are not shown. The pool of TACC3 that is not bound to clathrin or ch-TOG may be much larger in interphase cells. **(C)** Nanoscale model of a MT to show the conformation and approximate location of the two MT-resident pools. The complex of TACC3–ch-TOG binds to the distal tip of the MT. Recognition of the tip is by ch-TOG, which occurs in an autonomous manner. EB proteins can bind in a zone 30-60 nm away (Maurer et al., 2014; Zanic et al., 2013). Beyond this, TACC3–ch-TOG–clathrin complexes can bind. TACC3 is active in binding, but only in partnership with clathrin (Hood et al., 2013). The full clathrin triskelion and cross-bridges to other MTs are not shown for simplicity.

## Discussion

In this paper we have described how TACC3 and ch-TOG interact at the plus-ends of MTs in human cells in interphase or mitosis. The localization of TACC3 to the MT plus-end depends on ch-TOG and does not require EB1/3 proteins, nor does it require Aurora-A phosphorylation and clathrin binding. Although the +TIP behavior of TACC3 is clear, we could detect no change in microtubule dynamics as a result of modulating TACC3 levels in cells.

TACC3–ch-TOG track the very distal tips of MTs, ahead of EB proteins. We measured that on average, TACC3 is localized 229 nm ahead of EB3. Work in *Xenopus* neuronal growth cones placed xTACC3 500 nm ahead of EB1 (Nwagbara et al., 2014). Our measurement is closer to the 114 nm measured for ch-TOG in human cells by super resolution microscopy (Nakamura et al., 2012), and for XMAP215 on in vitro MTs by a superior fitting method that accounted for the PSF of the optical system (Maurer et al., 2014; Zanic et al., 2013). Our simplistic measurement is therefore likely to be an overestimate. All of the results are consistent with a model whereby TACC3–ch-TOG binds the distal tip of the MT where ch-TOG is involved in MT polymerization and EB proteins bind in a zone further away. The TACC3–ch-TOG–clathrin complex binds via a different mode that does not involve ch-TOG. This MT lattice-binding mode is occupies the rest of the MT, away from the EB zone and the distal tip (Fig 8C).

Our data show that TACC3 binds ch-TOG via an interaction between a break in the coiled-coil of the TACC domain and two leucine residues in the 4α domain of ch-TOG. Structural information from small angle X-ray scattering (SAXS) is now available for xTACC3 (Mortuza et al., 2014), which proposes that the TACC domain is a bundle of four alpha helices. The ch-TOG-binding region on TACC3 that we identified sits in α2 in the xTACC3 structural model, and mutations in this region abrogate binding to XMAP215 (Mortuza et al., 2014). Interestingly, they identified a different region at the extreme C-terminus of XMAP215 that interacts with xTACC3 (Mortuza et al., 2014). Note that their *in vitro* work used a 60-residue peptide, whereas we used full-length proteins in cells and longer, folded domains for our biochemical work. Nonetheless, it will be interesting to determine if multiple contacts between these two molecules are made.

The expression of TACC3 is altered in different human cancer types (Lauffart et al., 2005; Ma et al., 2003; Williams et al., 2013). Previous work explored the possibility that overexpression of TACC3 increases cell invasiveness and promotes a more aggressive cancer phenotype (Ha et al., 2013). The authors proposed that TACC3 promotes EMT via alteration of signaling pathways. We wondered if altered microtubule dynamics may explain the changes in cell migration. Our experiments confirmed that depletion of TACC3 reduces migration speed. However, we found that altering TACC3 expression during interphase did not detectably change MT dynamics. This was disappointing, because other +TIP proteins have been shown to be involved in cell migration, e.g. GTSE1 (Scolz et al., 2012) a protein reported to bind TACC3 (Hubner et al., 2010). An intriguing possibility is that the decrease in cell migration speed in TACC3-depleted cells is via a change in cell adhesion, since ENAH and VASP have previously been identified as TACC3 binding partners (Hubner et al., 2010).

What is the function of TACC3 as a +TIP? We could see no clear effect of TACC3 on MT dynamics, but it is possible that we have overlooked a role for TACC3 in modulating MT dynamics via TACC3–ch-TOG. For example, it was not possible for us to assess the function of TACC3 as a +TIP during mitosis, since TACC3 participates in a different function with ch-TOG and clathrin during this time. If the +TIP activity in mitosis is important for modulating MT growth it may be involved in acentrosomal MT assembly during prometaphase (Fu et al., 2013). The other recently described role for TACC3 in anaphase MT sliding (Lioutas and Vernos, 2013) is dependent on Aurora-A kinase activity and so likely involves the TACC3–ch-TOG–clathrin pool rather than the TACC3–ch-TOG +TIP activity. In *Xenopus* neuronal growth cones, only minor changes (~11 %) in MT dynamics were caused by xTACC3 depletion or overexpression (Nwagbara et al., 2014). This suggests that TACC3 may not significantly influence ch-TOG activity at plus-ends. However, a bigger effect was observed in neural crest cells, but only for TACC3 depletion (Nwagbara et al., 2014). Other groups have shown that TACC3 influences MT regrowth after depolymerization (Singh et al., 2014; Yang et al., 2012), although this may be unrelated to the +TIP function of TACC3. The observation of TACC3 piggybacking on ch-TOG at MT plus-ends is tantalizing, but firm evidence of a functional correlate of TACC3 +TIP behavior is lacking.

Figure 8 shows the subcellular pools of TACC3, ch-TOG and clathrin that can be detected alone and in combination in human cells. We do not place TACC3 at the centrosome in this scheme. TACC3 is often described as a centrosomal protein (Thakur et al., 2013). The evidence that TACC3 is present at centrosomes comes from two studies. First, TACC3 was observed at spindle poles in nocodazole-treated cells (Gergely et al., 2000a). However, this treatment leaves MT remnants that TACC3–ch-TOG–clathrin may bind to, rather than directly at the centrosome. Second, an antibody raised against TACC3 phosphorylated at S558 recognized centrosomes (Kinoshita et al., 2005). However, later work showed that this antibody simply detects centrosomes and not TACC3 (Lin et al., 2010). The signal for TACC3 pS558 detected with a new antibody was found over the K-fibers and not at the centrosomes (Lin et al., 2010). Although we note that a recent paper indicates that TACC3 may interact γ-tubulin complex components (Singh et al., 2014). One further reason that TACC3 is known as a centrosomal protein is historical. Initial work on TACC3 focused on potential similarities with the TACC homologs of lower species, where the relevant cell biology had originally been investigated, and in which TACC homologs are clearly found at the centrosomes or spindle pole body (Gergely et al., 2000a; Gergely et al., 2000b; Peset et al., 2005; Sato et al., 2004). More recently however, multiple studies including the present work have failed to show obvious localization of TACC3 at the centrosome in mitotic higher eukaryotic cells (Fu et al., 2010; Gergely et al., 2003; Hood et al., 2013; Lin et al., 2010). Since growing plus-ends of MTs continually emerge from the centrosome/spindle pole, this might explain apparent localization of TACC3 at the centrosome that may have been previously observed (Fig 1). Note that EB3 also has a “centrosomal” appearance, presumably for the same reason (Fig 1).

By contrast, it is clear that ch-TOG has a centrosomal distribution in interphase and mitosis (Fig 7) (Booth et al., 2011; Charrasse et al., 1998; Foraker et al., 2012; Gergely et al., 2003). This localization persists following knockdown of TACC3 or in the absence of an interaction with TACC3 (Fig 3) (Booth et al., 2011; Gergely et al., 2003; Lin et al., 2010). By contrast, in *Drosophila*, D-TACC – the sole TACC3 homolog – is required to localize mini spindles (msps) to centrosomes (Lee et al., 2001). In fly oocyte spindles D-TACC localizes msps to acentriolar poles via kinesin-14 transport (Cullen and Ohkura, 2001). The kinetochore localization of ch-TOG has not been reported previously although Alp14, an *S. pombe* homolog of ch-TOG, localizes at kinetochores (Garcia et al., 2001; Sato et al., 2004) as does Msps, in *Drosophila* (Buster et al., 2007). This pool of ch-TOG is not readily detected by immunofluorescence (Booth et al., 2011; Charrasse et al., 1998; Lin et al., 2010), suggesting that it is inaccessible to antibodies directed against the C-terminus of ch-TOG. It is unclear why TACC3 is unable to bind the ch-TOG that is localized at centrosomes and kinetochores in human cells. It could be that TACC3 is prevented from doing so because ch-TOG is in a different conformation or that it is bound to another protein. These are interesting questions for future study.

The scheme of TACC3, ch-TOG and clathrin pools that we describe here (Fig 8) represents a current, consolidated view of these fascinating proteins. It highlights how the switch from three pools (in interphase) to four (in mitosis) is achieved by a simple phosphorylation event mediated by Aurora-A kinase. We hope this will be a useful framework for interpreting and understanding future research in this area.

## Acknowledgements

We thank Rachel Jones for technical assistance, Anne Straube (Warwick Medical School) and Tony Hyman (MPI Dresden) for important reagents, Fanni Gergely (Cancer Research UK Cambridge Institute) for help in designing antigen peptides for the production of the TACC3 TACC domain antibody and Mark Richards (University of Leicester; Cancer Research UK Leicester Centre) for assistance with CD spectroscopy. Colleagues at Warwick and Leicester provided important critical discussion.

## Author contributions

CGC carried out all of the experimental work except for biochemical experiments in Figure 4, which were performed by SGB. SJR analyzed data and wrote the paper. All authors injected ideas and criticism, contributed to the editing of the manuscript and approved the final version.

## Supplementary Information

### Supplementary Movie 1 A video of GFP-TACC3 expressed in an interphase RPE1 cell stably expressing EB1

Live-cell imaging was done using a spinning disk microscope with a 100X ~1.4 NA oil immersion objective lens. Images were captured every second for 1 minute (video playback is 10 fps). Cells were excited at 488 nm and 561 nm and images captured simultaneously with two cameras (Hamamatsu C10600-10B ORCA-R2). The same cell is shown in Figure 2A.

### Supplementary Movie 2 A video of GFP-TACC3 expressed in a mitotic RPE1 cell stably expressing EB1

Live-cell imaging was done using a spinning disk microscope with a 100X ~1.4 NA oil immersion objective lens. Images were captured every second for 1 minute (video playback is 10 fps). Cells were excited at 488 nm and 561 nm and images captured simultaneously with two cameras (Hamamatsu C10600-10B ORCA-R2). The same cell is shown in Fig 1B.

### Supplementary Movie 3 A video of a HeLa Kyoto cell in anaphase expressing TACC3-GFP under its endogenous promoter

Live-cell imaging was done using a spinning disk microscope with a 100X ~1.4 NA oil immersion objective lens. Images were captured every second for 1 minute (video playback is 10 fps). The same cell is shown in Fig 1C.

